# Amyloid particles facilitate surface-catalyzed cross-seeding by acting as promiscuous nanoparticles

**DOI:** 10.1101/2020.09.01.278481

**Authors:** Nadejda Koloteva-Levine, Ricardo Marchante, Tracey J. Purton, Jennifer R. Hiscock, Mick F. Tuite, Wei-Feng Xue

## Abstract

Amyloid seeds are nanometre-sized protein particles that accelerate amyloid assembly, as well as propagate and transmit the amyloid protein conformation associated with a wide range of protein misfolding diseases. However, seeded amyloid growth through templated elongation at fibril ends cannot explain the full range of molecular behaviours observed during cross-seeded formation of amyloid by heterologous seeds. Here, we demonstrate that amyloid seeds can accelerate amyloid formation via a surface catalysis mechanism without propagating the specific amyloid conformation associated with the seeds. This type of seeding mechanism is demonstrated through quantitative characterisation of the cross-seeded assembly reactions involving two non-homologous and unrelated proteins: the human Aβ42 peptide and the yeast prion-forming protein Sup35NM. Our results suggest experimental approaches to differentiate seeding by templated elongation from non-templated amyloid seeding, and rationalise the molecular mechanism of the cross-seeding phenomenon as a manifestation of the aberrant surface activities presented by amyloid seeds as nanoparticles.

## INTRODUCTION

Amyloid particles are associated with numerous neurodegenerative and/or age-related human disease such as Alzheimer’s disease, Huntington’s disease, Parkinson’s disease and type 2 diabetes mellitus (Eisenberg and Jucker, 2012; Knowles et al., 2014). The slow nucleation-dependent process that converts normally soluble protein or peptide precursors into their amyloid conformation (Xue et al., 2008) can be bypassed through the addition of preformed amyloid particles, the seeds (**Figure 1**). This phenomenon, which effectively accelerates amyloid growth and propagates the amyloid conformation, is called seeding. The seeded growth of amyloid, as well as transmissible forms of amyloid known as prions, via templated the addition of monomers or small oligomers to the ends of pre-formed fibril seeds (Collins et al., 2004; Ferrone, 1999; Xue, 2015), is known as fibril elongation.

**Figure 1.**
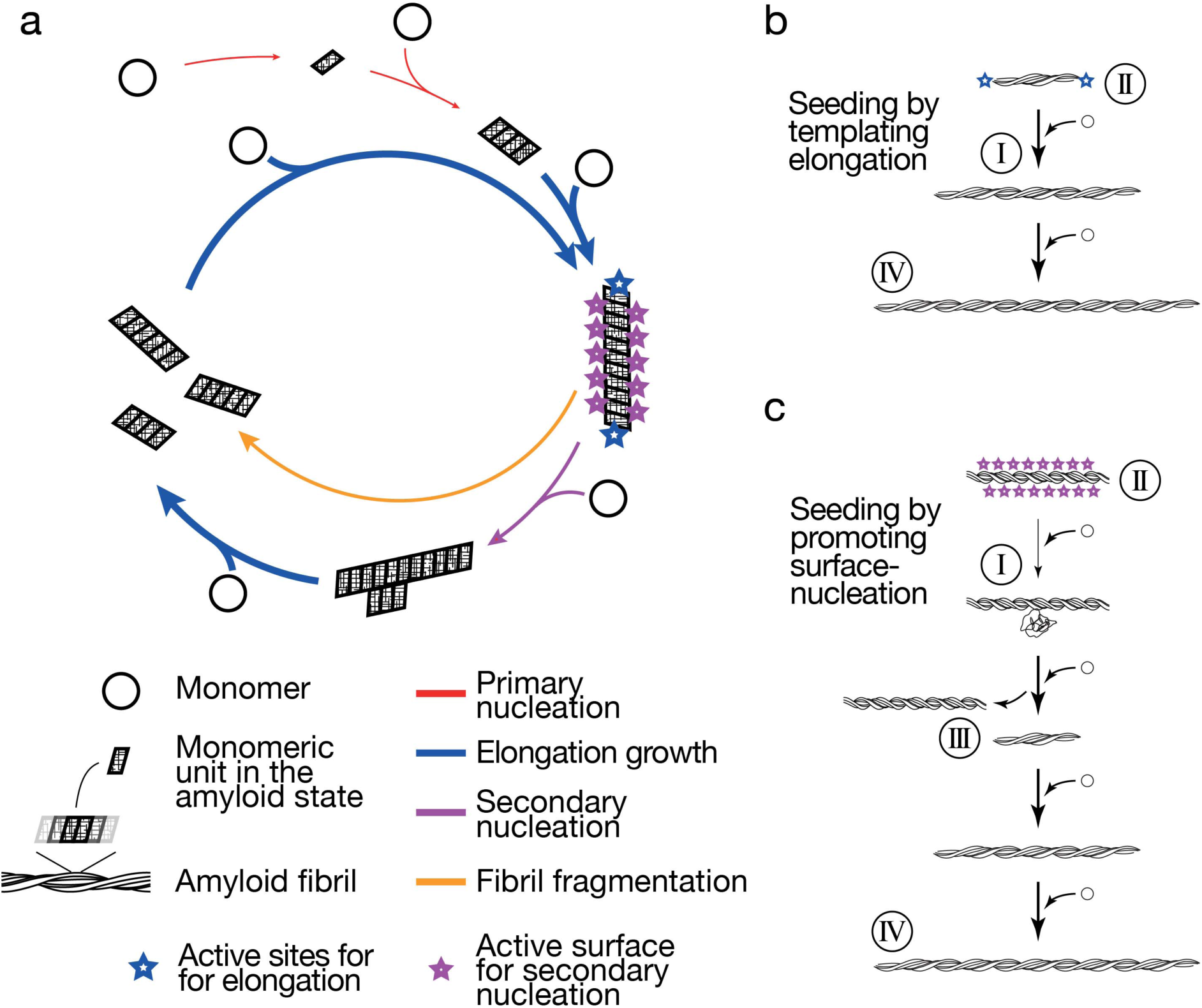
Schematic illustration of the molecular processes in the amyloid life-cycle together with experimentally testable hypotheses. The amyloid life-cycle (a) with key sites and surfaces for templated growth and secondary nucleation highlighted, respectively. (b) Schematic illustration of a seeding mechanism based on template growth at fibril ends. (c) Schematic illustration of a seeding reaction with fibril growth promoted by a fibril surface-catalyzed nucleation mechanism. Numbers indicate experimentally testable and comparable features of the mechanisms. I) Prediction: A surface-catalyzed seeding reaction is nucleation dependent and therefore slow compared to seeding through elongation. II) Prediction: The number of active sites (blue stars) for seeding through templated elongation depends on the number of particles, while the number of active surface for surface-catalyzed seeding mechanism depends on the protein mass concentration. III) Prediction: The morphology of the fibrils newly formed by surface-catalyzed seeding does not need to be the same as that of the seeds. IV) Prediction: Indistinguishable fibril morphology and biological activity is produced from the same monomers under the same conditions independently of the seeds used for a surface-catalyzed seeding reaction. The arrows represent dynamic and reversible steps along the lifecycle and the thickness of the arrows illustrate typical relative magnitudes of the rates involved in each of the processes.

Elongation at amyloid fibril ends has long been viewed as the sole mechanism of seeded amyloid growth, where the specific amyloid conformation encoded in the seeds is propagated upon the addition of new monomers or small oligomers at fibril ends (Collins et al., 2004; Serio et al., 2000). This mechanistic assumption has been, for example, applied in propagating patient-derived amyloid material for structural studies. However, a protein can form amyloid in an accelerated manner upon addition of amyloid seeds preformed with precursors of very different or even completely non-homologous amino-acid sequences (Sarell et al., 2013). The molecular mechanism of this phenomenon, often termed ‘cross-seeding’, remains unresolved because the current models for the fibril elongation growth mechanism cannot explain the full range of molecular behaviours observed during amyloid cross-seeding.

For mammalian disease-associated amyloidogenic proteins, cross-seeding activity may be a key process promoting a synergy between amyloid associated disorders. A number of studies have demonstrated that two different amyloidogenic disorders may arise in the same individual and, in so doing, impact on the respective occurrence and pathologies of the disorders (Lundmark et al., 2005; Morales et al., 2013). For example, it has been proposed that cross-seeding between Aβ and α-synuclein (Ono et al., 2012), and Aβ_42_ and IAPP (O’Nuallain et al., 2004) might contribute the observed statistical correlations between the occurrence of Alzheimer’s disease, and Parkinson’s disease or type 2 diabetes, respectively (Biessels et al., 2006; Daniele et al., 2018; Sims-Robinson et al., 2010). Thus, understanding the fundamental nature of the molecular cross-talk between amyloidogenic disease-associated proteins and the underlying molecular mechanism will provide essential clues to a better understanding of how these diseases originate, propagate and even transmit between individuals.

Amyloid fibrils are protein filaments with monomeric units arranged in the characteristic cross-beta molecular architecture held together by non-covalent interactions and hydrogen bonds parallel to the fibril axis (Sunde et al., 1997). The fibrils are usually in the order of 10 nm in width, and amyloid seeds are typically small amyloid fibril fragments often less than 100 nm in length. Thus, amyloid seeds are *bona fide* nanoparticles, i.e. particulate materials with individual particle dimensions in the order of or below 100 nm for at least two out of three spatial directions (Vert et al., 2012). Like any type of nanoparticle, the small sizes of amyloid seeds confer these particles with high surface-to-volume ratios. These surfaces are frequently capable of interacting with molecules in their surroundings, and can be coated with a corona of proteins and other macromolecules in addition to ions and small molecules in biological environments (Cedervall et al., 2007; John et al., 2018).

The surfaces of amyloid seeds contain active sites where growth by elongation takes place at their termini, as well as surfaces parallel to the cross-beta hydrogen bonds that have previously been shown to be exceptionally active in catalyzing heterogeneous nucleation of new amyloid in what is called ‘secondary nucleation’ as is observed in the formation of Aβ amyloid (Cohen et al., 2013). Thus, amyloid seeds, in addition to promoting the propagation of the specific amyloid conformation encoded by the monomeric units at their elongation active sites through templated growth at fibril ends, may also be able to catalyze generic surface mediated assembly like any nanoparticle, and accelerate the formation of heterologous amyloid through catalyzing heterogeneous nucleation.

In order to test whether such a general surface-catalyzed mechanism can explain and rationalize the molecular mechanism of amyloid cross-seeding, and to show that the seeding and templating activities of amyloid seeds can potentially be mechanistically uncoupled, we investigated the cross-seeding interactions between two unrelated amyloidogenic proteins: human Aβ42 that is associated with Alzheimer disease (Knowles et al., 2014) and the amyloid-forming protein Sup35NM that is a component of a prion-based epigenetic switch in the yeast *Saccharomyces cerevisiae* (Serio et al., 2000). We have chosen these two proteins because being from two organisms at different ends of the evolutionary spectrum, they do not co-exist in the same biological context. Sup35 (eRF3) is present in human cells but lacks the NM region critical for amyloid formation and propagation (Serio et al., 2000). Furthermore, Aβ and Sup35NM have dissimilar amino acid compositions and low sequence similarities (**Supplementary Figure SI-1**) as expected from two functionally-unrelated proteins. The amyloid aggregation mechanism of human Aβ42 has been studied in considerable detail (Cohen et al., 2013), revealing an assembly mechanism dominated by secondary nucleation accelerated by pre-existing fibril surfaces. Sup35, on the other hand, is regarded as a functional amyloid, and its assembly mechanism reveals a strong component of templated elongation (Collins et al., 2004; Serio et al., 2000). Here we demonstrate that these two unrelated proteins are capable of cross-seeding each other, i.e. the presence of the amyloid seed of one protein is capable of accelerating amyloid formation of the other protein despite their dissimilar sequence, structure and biological origins. We also show that these heterologous interactions are ‘asymmetric’ (Kumar and Udgaonkar, 2018), i.e. the kinetic effect of the seeds are different with respect to amyloid assembly of each other. We demonstrate that these cross-seeding interactions are mass sensitive, but not particle number sensitive. In addition, by exploiting the well characterized prion phenotype [*PSI*^+^] associated with the amyloid state of Sup35 protein, we demonstrate the phenotypic outcome on cells propagating either the self-seeded or cross-seeded Sup35NM prion particles *in vivo*. Together, our *in vitro* and *in vivo* results demonstrate that amyloid seeds are nanoparticles with active surfaces that are capable of mediating the cross-seeding of heterologous amyloid through generic surface-catalyzed reactions, resulting in accelerated amyloid growth, without structural templating of the precise amyloid conformation encoded in the seeds.

## RESULTS

### Both templated elongation and surface-catalyzed nucleation mechanisms can promote accelerated amyloid assembly but each produce different seeding behaviours

Seeding is a process defined as the acceleration of amyloid formation in the presence of seeds. In order to test whether amyloid seeds are nanoparticles that can accelerate the formation of new heterologous amyloid through non-templated surface-catalyzed assembly reactions, we first examined in detail the mechanistic differences between templated seeding reactions by elongation versus the non-templated surface-catalyzed seeding reactions (**Figure 1b compared to Figure 1c**). In so doing we sought to establish whether the addition of seeds can accelerate amyloid assembly either through templated growth by elongation at fibril ends (**Figure 1b**) or surface-catalyzed nucleation of new amyloid (**Figure 1c**), and if these two pathways can be distinguished experimentally. As illustrated in **Figure 1**, if given amyloid particles are capable of seeding or cross-seeding the formation of new amyloid through a surface-catalyzed nucleation mechanism instead of templated elongation, one would predict a number of differences in the molecular and kinetic behaviours of the seeded reactions that should be distinguishable experimentally.

Firstly, the presence of active surfaces should reduce but not eliminate the nucleation barrier for assembly (**Figure 1, prediction I**), since surface-catalyzed assembly would still be a nucleation-dependent process. Consequently, such a reaction would still go through a slow nucleation phase, albeit faster compared to nucleation in the absence of surface catalysis. Thus, the addition of seeds active in surface catalysis of nucleation would only be capable of reducing the length of the lag phase, but not eliminate the lag phase entirely as would be observed with seeding reactions that proceed through templated elongation. Secondly, the number of growth active sites for templated elongation is only present at fibril ends, and therefore relates to the particle concentration of the seed particles. On the other hand, surfaces along the seeds that potentially can catalyze heterogeneous nucleation such as secondary nucleation or surface-catalysed seeding events should depend on the total length of the particles, which is in turn proportional to the mass or monomer equivalent concentration of seeds, and not the number concentration of the seed particles (**Figure 1, prediction II**). Thirdly, if new amyloid is formed through seeding by a surface-catalysis mechanism then the fibril morphology of the newly formed fibrils does not need to be the same as the morphology of the seeds (**Figure 1, prediction III**). Finally, if newly formed amyloid assembles through seed surface-catalyzed reactions then their morphology and the biological response they elicit should only be linked to their monomer precursors and the conditions applied but not the seeds (**Figure 1, prediction IV**). These four experimentally testable differences were therefore used to rationalize whether amyloid particles can act as broad spectrum seeds that are capable of accelerating the formation of new and heterologous amyloid primarily due to the activities of their surfaces in the same way as the action of nanoparticles.

### Self-seeded growth of both Aβ42 and Sup35NM amyloid fibrils proceed through templated fibril elongation

In order to characterize the heterologous seeding potential of amyloid particles, we chose to investigate the self-seeding and cross-seeding interactions between two unrelated proteins: human Aβ42 and yeast Sup35NM. These two amyloid-forming proteins are native to different organisms, do not naturally co-exist in the same biological context, and do not share any known evolutionary linkages. Furthermore, human Aβ is associated with Alzheimer’s disease known to be statistically correlated with the occurrence of a number of other amyloid associated diseases (Biessels et al., 2006; Daniele et al., 2018; Sims-Robinson et al., 2010), while the full-length Sup35 protein can become a transmissible prion and can be regarded as a functional amyloid (Serio et al., 2000). These two proteins also have low sequence similarities (**Supplementary Figure SI-1c**), and are dissimilar in terms of size, charge, amino acid composition (**Supplementary Figure SI-1**) and fibril structures. Therefore, each of these two proteins should not be able to grow onto fibril seeds pre-formed by the other protein through templated assembly. Hence, they provide ideal and unbiased tests of the heterologous seeding capabilities of amyloid particles.

We first generated fibrillar particles of Aβ42 and Sup35NM *in vitro* by incubating monomeric precursors under common fibril growth solution conditions as both proteins form amyloid fibrils under physiological pH. To generate Aβ42 fibril particles, a synthetic Aβ42 peptide (Bachem, Germany) was used. The peptide samples were dissolved in 6M GdnHCl at pH10 and Aβ42 monomers were purified by gel filtration using a Superdex 75 column immediately prior to assembly in order to ensure the generation of reproducible and high quality Aβ42 amyloid seeds (Hellstrand et al., 2010; Walsh et al., 2009). Monomeric Sup35NM protein was produced recombinantly in *E. coli* and assembled as described previously (Marchante et al., 2017). Aβ42 and Sup35NM fibrils were subsequently dispersed by brief controlled sonication (see Materials and Methods) and imaged using atomic force microscopy (AFM). Interestingly, the Aβ42 particles were apparently less resistant to sonication compared to Sup35NM particles. Consequently, 1s of controlled sonication was sufficient to disperse the Aβ42 fibrils for imaging, while at least 5s of controlled sonication was required to disperse Sup35NM fibrils prior to imaging. **Figure 2a** shows AFM images of Aβ42 and Sup35NM fibril seeds imaged by AFM.

**Figure 2.**
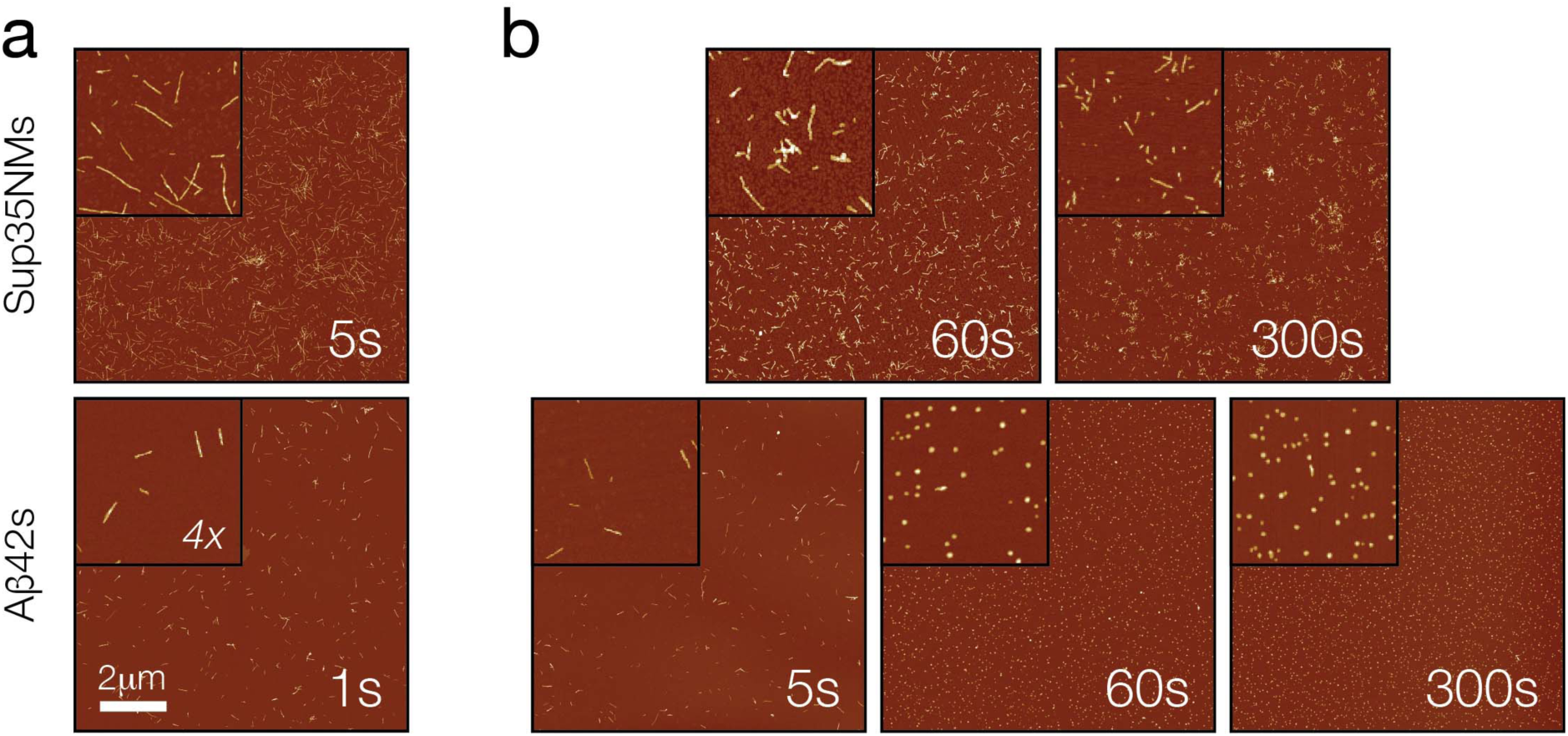
AFM images of Aβ42 and Sup35NM fibril seeds. (a) AFM images of Aβ42 and Sup35NM seeds after brief controlled sonication to disperse the fibrils. (b) AFM images of Aβ42 and Sup35NM seeds after different length of controlled sonication.

Controlled sonication is a method commonly used to fragment amyloid fibrils for seed generation. To validate controlled sonication as being capable of generating Aβ42 and Sup35NM seed samples with different particle concentrations, while retaining their respective original mass concentrations, the seed samples were subjected to controlled sonication for different periods of time. As seen in **Figure 2b**, for Sup35NM, increasing the sonication time from 5 s to 300 s significantly decreased the lengths of the seed particles, and therefore, increased the particle concentration as expected and previously seen (Marchante et al., 2017). The same effect of decay in particle lengths and a rise in particle concentration as sonication time increased from 1 s to 60 s was also seen for Aβ42 particles (**Figure 2b**). As previously observed, the Aβ42 particles were less resistant to sonication compared to Sup35NM particles, with 60 s of controlled sonication generating a large number of small nanoparticles less than 100 nm as seen using AFM (**Figure 2b**). Further sonication did not further alter their size distribution noticeably as would be expected due to their already small sizes (Beal et al., 2020; Xue and Radford, 2013). To further confirm the quality the Aβ42 and Sup35NM seed samples, dynamic light scattering (DLS) was performed on these seed samples after controlled sonication (Materials and methods). As shown in **Supplementary Figure SI-2**, the DLS experiments show that both the Aβ42 and Sup35NM seed samples consisted of a distinct distribution of fragmented fibrils without the presence of any major secondary particle distributions. Thus, the AFM and DLS experiments together confirmed the presence of high-quality seed samples for both Aβ42 and Sup35NM formed under the same solution conditions.

To confirm the ability of Aβ42 and Sup35NM particles to seed the formation of new amyloid, we performed a series of seeded fibril growth kinetic assays monitored using the fluorescence of the amyloid-specific Thioflavin T dye in a 96-well plate format, involving both Aβ42 and Sup35NM each seeded by seeds formed from monomers of the same sequence (i.e. self-seeding). The kinetic profiles of amyloid growth were mapped as a function of low seed concentrations ranging from 0.1 to 5% to determine the parameters of respective seeded growth and their dependence on seed particle concentration (**Figure 3a and 3d**). Subsequently, we extracted and analysed two parameters characteristic for the amyloid assembly kinetics (**Supplementary Figure SI-3**); the length of the lag phase (*t*_*lag*_, **Figure 4**) and the initial slope (*k*_*0*_, **Figure 5**) from each reaction trace. For Sup35NM fibril seeds (Sup35NMs) self-seeded with Sup35NM monomers (Sup35NMm), the presence of 0.1-5% monomer molar equivalent seeds in a growth reaction with 10 µM total monomer equivalent concentration dramatically shortened or eliminated the lag-phase in all cases. Importantly, as seen in **Figure 4**, as little as 0.5-1% monomer molar equivalent of the seeds was sufficient to completely eliminate the lag phase by reducing *t*_*lag*_ to 0 h. This behaviour is entirely consistent with templated monomer addition to the pre-formed fibril-seed ends acting as a dominant mechanism of the elongation growth (Collins et al., 2004; Serio et al., 2000). Seeding efficiency also increased with higher added particle concentrations either through increased monomer molar equivalent seeds or through increased sonication at the same monomer molar equivalent (**Figure 4**). As seen in **Figure 5**, the initial slope of seeded growth curves is directly proportional to particle concentration, which in turn is proportional to the number of active growth sites for elongation at fibril-ends, at low particle concentrations. At high concentrations, the elongation process can become saturated as also seen in other amyloid systems (e.g. (Buell et al., 2014)). Therefore, the data demonstrate that the self-seeded growth of Sup35NM proceeds through templated elongation at fibril ends as expected (Collins et al., 2004).

**Figure 3.**
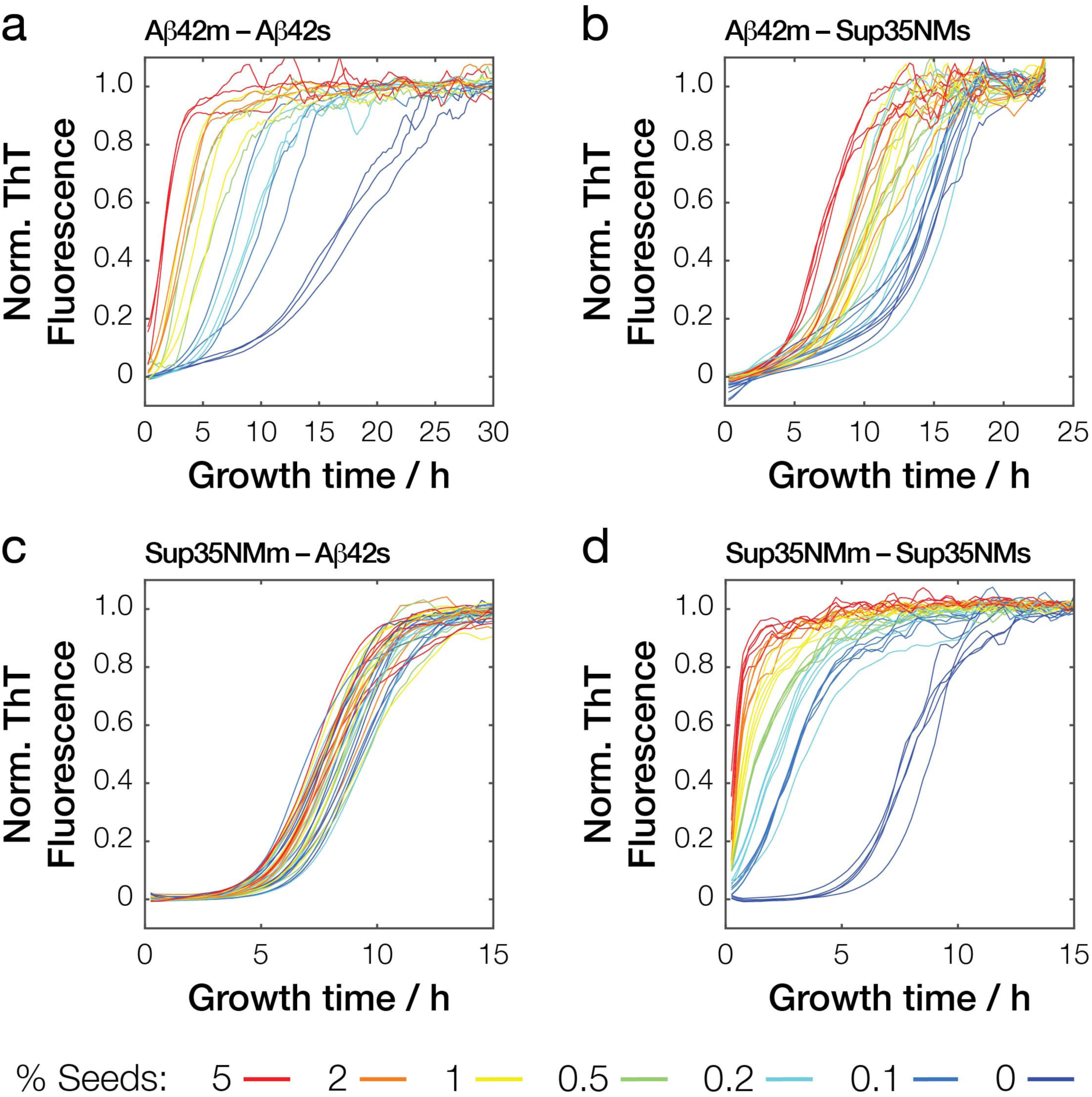
Kinetics traces of seeded amyloid formation monitored by ThT fluorescence. Typical normalised traces of (a) monomers of Aβ42 (Aβ42m) self-seeded by Aβ42s or (b) by Sup35NM seeds, as well as (c) Sup35NM monomers (Sup35NMm) seeded by Aβ42s or (d) self-seeded by Sup35NMs seeds. For each, reactions with difference ratios (0-5%) of seeds added are shown.

**Figure 4.**
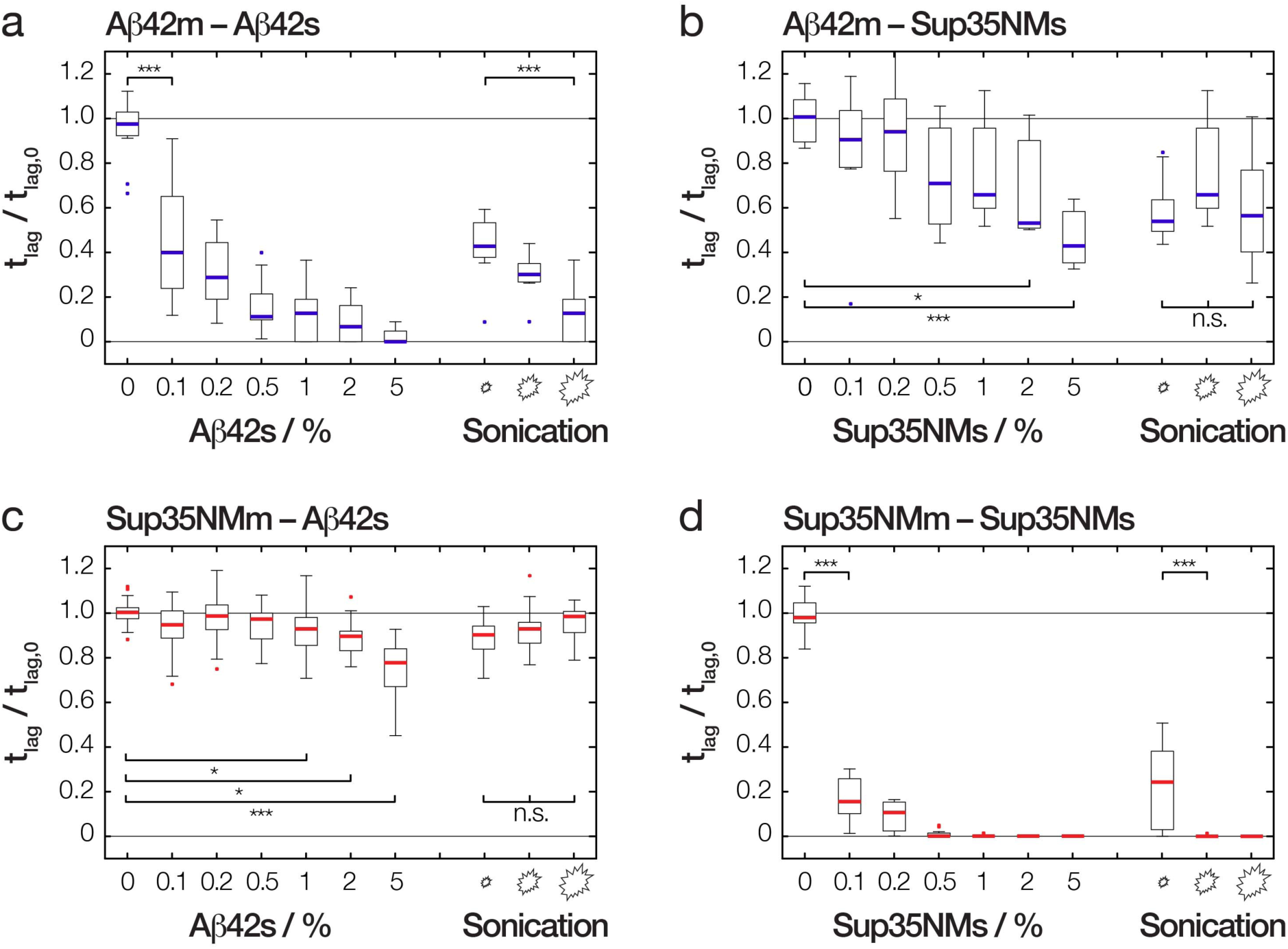
The relative reduction in the length of lag phase upon addition of seeds compared to unseeded amyloid formation. The relative *t*_*lag*_ values of (a) Aβ42 (Aβ42m) self-seeded by Aβ42s or (b) by Sup35NM seeds, as well as (c) Sup35NM monomers (Sup35NMm) seeded by Aβ42s or (d) self-seeded by Sup35NMs seeds are shown as ratio to *t*_*lag*_ values of respective unseeded reactions (*t*_*lag,0*_). For each protein pair, seeding reactions were performed with varying % seeds as well as varying degree of sonication as indicated. The *t*_*lag*_ values were extracted from the kinetics traces using the method illustrated in **Supplementary Figure SI-3**. The distribution of *t*_*lag*_ values for each experiment are shown as box plots with the thick line representing the median, and each bar represents the data from at least 9 replicate reactions from 3 independent experiments.

**Figure 5.**
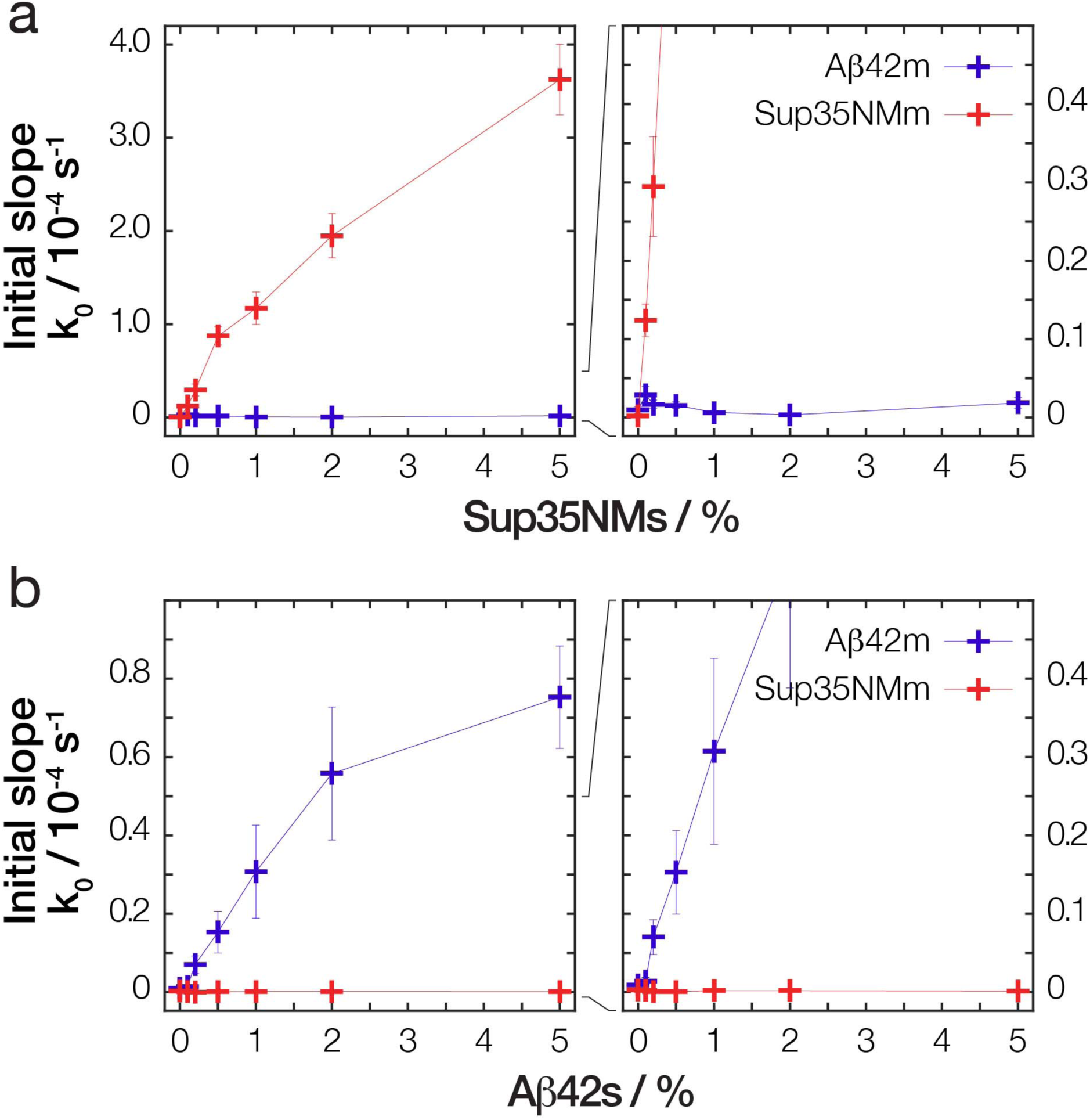
The increase of the initial slope of the kinetic traces as function of increasing percentage of seeds. The average initial slope values (*k*_*0*_) from traces of (a) Aβ42 (Aβ42m) or Sup35NM (Sup35NMm) seeded by Sup35NM seeds, as well as (b) seeded by Aβ42s are shown as ‘+’ with error bars representing the standard error of mean. The *k*_*0*_ values were extracted from the kinetics traces using the method illustrated in **Supplementary Figure SI-3**. The error bars for some reactions with low *k*_*0*_ values are not visible due to being smaller than the symbol.

Similarly to Sup35NM, assembly of Aβ42 monomers (Aβ42m) was significantly accelerated by the addition of as little as 0.1% pre-formed Aβ42 fibril-seeds (Aβ42s). The lag-phase was eliminated in the presence of as low as ∼1% monomer molar equivalents or more of seeds (**Figure 4**). Increasing the seed particle concentration also increased the initial slope of the growth reactions in the same linear manner at low seed particle concentrations as seen with Sup35NM and other amyloid-forming systems (**Figure 5**). Furthermore, increasing the particle concentration at constant molar equivalent monomer concentration of seeds by sonication also shortened the lag-phase, demonstrating that the seeding reaction is dependent on the particle concentration of the seeds. Therefore, the self-seeded growth of Aβ42 amyloid also display all the hallmarks of a seeding mechanism dominated by templated elongation at fibril-seed ends.

### Growth of Aβ42 amyloid fibrils is accelerated by Sup35NM seeds in a mass concentration dependent, but not particle number dependent, manner

Having established the self-seeded reactions proceed through templated elongation at fibril-seed ends for both Aβ42 and Sup35NM, we next investigated whether the seeds formed from these two unrelated amyloidogenic proteins are able to accelerate the amyloid forming reaction of each other in cross-seeded reactions. First, we investigated the growth kinetics of Aβ42m assembly in the presence of Sup35NMs seeds. In a series of fibril growth kinetic assays monitored using Thioflavin T fluorescence (**Figure 3b**), the addition of heterologous seeds (in this case the unrelated Sup35NMs) in low concentrations (∼5% or less monomer molar equivalents) to monomer solutions of Aβ42 was able to statistically significantly reduce the lag time of the Aβ42 amyloid formation (**Figure 4b**). These experiments show that Sup35NMs is capable of acting as seeds that accelerate amyloid formation of Aβ42, albeit with less efficiency in reducing the duration of the lag phase than Aβ42s. Interestingly, the addition of 5% monomer molar equivalents of Sup35NMs failed to eliminate the lag phase of Aβ42 amyloid growth (*t*_*lag*_ > 0 in **Figure 4b**, and *k*_*0*_ ≈ 0 in **Figure 5a**). This finding is not consistent with a templated elongation mechanism, but is consistent with Sup35NMs being nanoparticles that act as generic seeds through surface-catalyzed heterogeneous nucleation because the slow nucleation events should still occur (**Figure 1 prediction I**). If the nucleation events catalyzed by the large available surface area of Sup35NMs (and not active growth sites at fibril-ends along) directs the shortening of the lag-phase in these heterologous cross-seeded reactions, then the number of free ends that are responsible for the templating process should exert no significant effect on the efficiency of seeding (**Figure 1 prediction II**) as long as the monomer equivalent concentration (equivalent to the mass concentration of seeds) is maintained. We tested this prediction by adding Sup35NMs sonicated to different extents, and therefore should have identical monomer mass concentration but different number of seed particle concentrations (Marchante et al., 2017), to the Aβ42m solutions (**Figure 2b**). Remarkably, adding an identical mass of Sup35NM seeds that were subjected to less time of sonication did not significantly increase the length of lag-phase of Aβ42 assembly, nor did the length of lag phase decrease significantly when identical mass of Sup35NM seeds, which were subjected to more time of sonication, were added. Thus, changing the particle concentration of Sup35NMs while maintaining identical protein mass concentration of seeds produced no significant effects on the length of lag-phase of Aβ42 fibril forming reactions (**Figure 4b**). These results provided kinetic evidence to support that Sup35NMs, while biologically and structurally unrelated to Aβ42, are able to accelerate the amyloid formation of Aβ42 by acting as nanoparticles that provide their surface for catalyzing Aβ42 amyloid formation.

### Growth of Sup35NM amyloid fibrils is also accelerated by Aβ42 seeds, but not to the same extent

We next tested if Aβ42s were also able to accelerate the amyloid assembly of Sup35NMm in the opposite heterologous cross-seeded reaction. As before, we incubated Sup35NMm solutions in the presence of various amounts of Aβ42s and monitored the fibril growth kinetic using Thioflavin T fluorescence (**Figure 3c**). For Sup35NM amyloid formation, Aβ42s was also able to significantly shorten, but not eliminate, the length of the lag phase. However, this lag phase-shortening effect was much less in magnitude compared with the other three reaction pairs analysed, with around a 25% reduction in the length of the lag phase upon addition of 5% monomer molar equivalents of Aβ42s compared to unseeded reactions. (**Figure 4c** and **Figure 5b**). Similarly to the effect of Sup35s on Aβ42m assembly, a change in particle concentration without a change in monomer equivalent mass concentration of Aβ42s by varying sonication time for Aβ42s, led to no significant effect on the length of the lag phase of Sup35NMm assembly. These experiments indicate that Aβ42s particle surfaces are also able to accelerate the nucleation of Sup35NM amyloid fibrils, but the efficiency is less compared to the effect of Sup35NMs surfaces on Aβ42m assembly (compare **Figure 4b and 4c**). This asymmetry is consistent with the fact that the efficiency of surface-catalyzed heterogeneous nucleation mechanism for cross-seeding is dependent on the physical and chemical properties of the seed surfaces in the same way as any nanoparticle interaction with biology.

### Sup35NM formed through self-seeding and cross-seeding with Aβ42 seeds display indistinguishable fibril morphology and induce identical prion phenotypes *in vivo*

If the mechanism of heterologous cross-seeding involves generic seeding through heterogeneous nucleation catalyzed by surfaces of seeds as nanoparticles then the amyloid fibril morphology of the newly grown amyloid does not need to be the same as that of the seeds (**Figure 1 prediction III**). The morphology and the biological properties of the newly grown amyloid may also be the same under the same solution conditions and independently of whether they were formed through homologous self-seeding or heterologous cross-seeding (**Figure 1 prediction IV**). To test these structural predictions, we used AFM to analyze Sup35NM fibrils formed in two seeded reactions, either self-seeded with pre-formed Sup35NMs fibrils or cross-seeded with Aβ42s fibrils (**Figure 6a**). The morphology of Sup35NMm fibril grown by cross-seeding with Aβ42s (**Figure 6a**), characterised by the height distribution (**Figure 6b**), was strikingly different compared to that of Aβ42s. The fibril heights of Aβ42s seeded Sup35NM fibrils were significantly different to that of the Aβ42s, but indistinguishable to self-seeded or *de novo* grown Sup35NM fibrils. This is consistent with the specific amyloid conformation of Aβ42 seeds not being imposed on the newly formed Sup35NM fibrils, confirming the structural prediction III (**Figure 1 prediction III**). Importantly, because fibrils in these heterologous cross-seeded reactions were indistinguishable from those formed from self-seeding with Sup35NMs or from *de novo* assembly of Sup35NMm (**Figure 6a**), and the height distributions for all of these samples were also not significantly different from each other (**Figure 6b**), these fibril populations must be formed mainly due to the monomer sequence and the solution condition, in agreement with the structural prediction IV (**Figure 1 prediction IV**). Taken together, these results support the hypothesis that the heterologous seeds in this case merely accelerated the kinetics of amyloid formation without passing on its precise conformation by a lack of templating.

**Figure 6.**
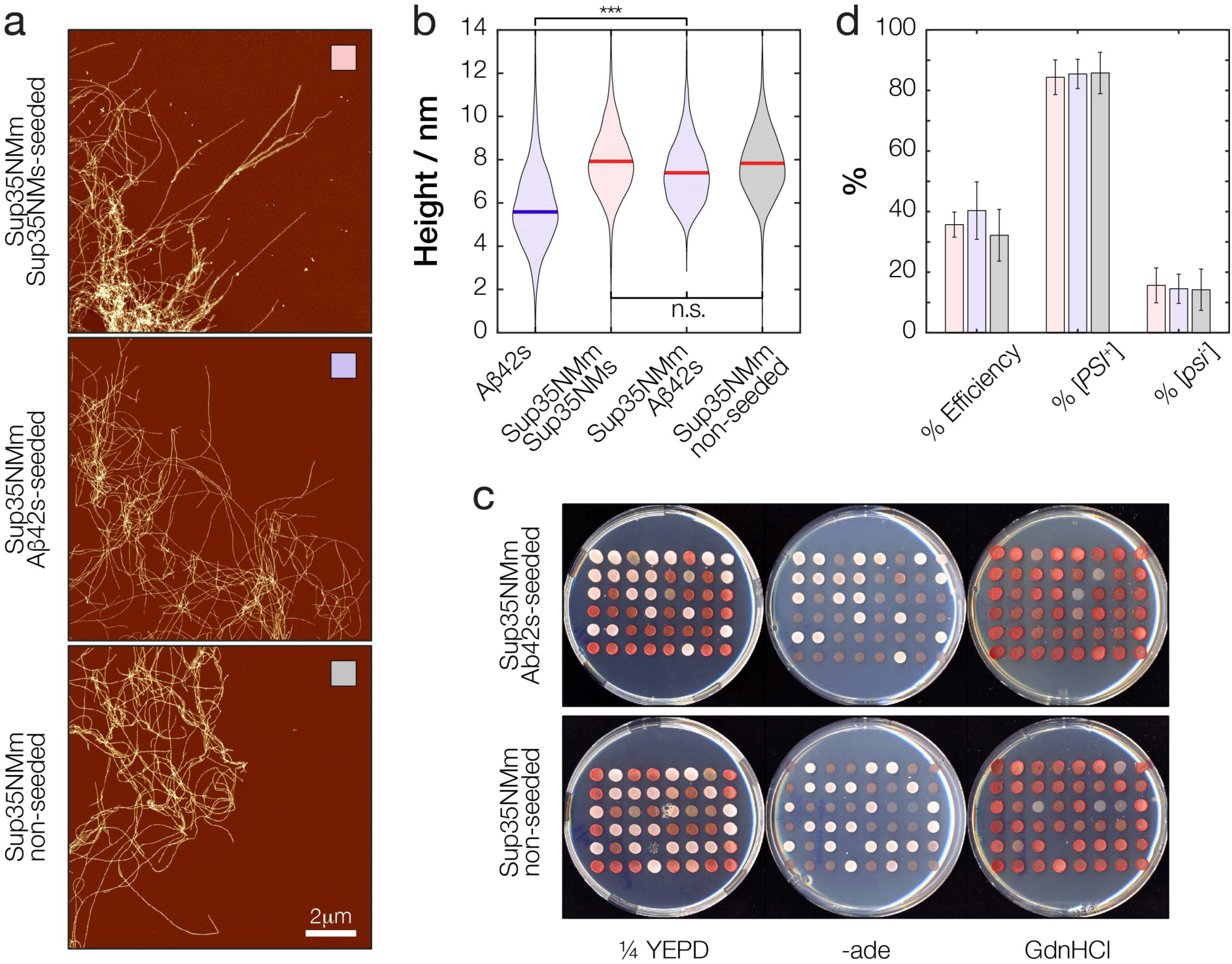
Sup35NM amyloid fibrils formed through self-seeding, cross-seeding using Aβ42s, or unseeded reactions are morphologically and biologically indistinguishable. (a) Typical AFM images of Sup35NM fibrils formed through self-seeding (upper), cross-seeding using Aβ42s (middle) or unseeded (lower) reactions. (b) The height distribution of the Sup35NM fibrils extracted from the images in (a) represented as violin plots. The thick line in each distribution represents the mean height. (c) Yeast cells transfected with the fibrils shown in (a) and then replica plated onto ¼ YEPD and -ade synthetic media to check for the [*PSI*^*+*^] prion phenotype and ¼ YEPD supplemented with 3mM GdnHCl to eliminate any false positives. The fibrils were sonicated for 600 s before transfection experiments. (d) Quantification and comparison of the transfection efficiency and the [*PSI*^*+*^] phenotype displayed by the yeast cells transfected with the fibrils shown in (a). The bars indicate average values of at least three independent experiments performed on separate days and the error bars represent the standard error of mean.

**Figure 7.**
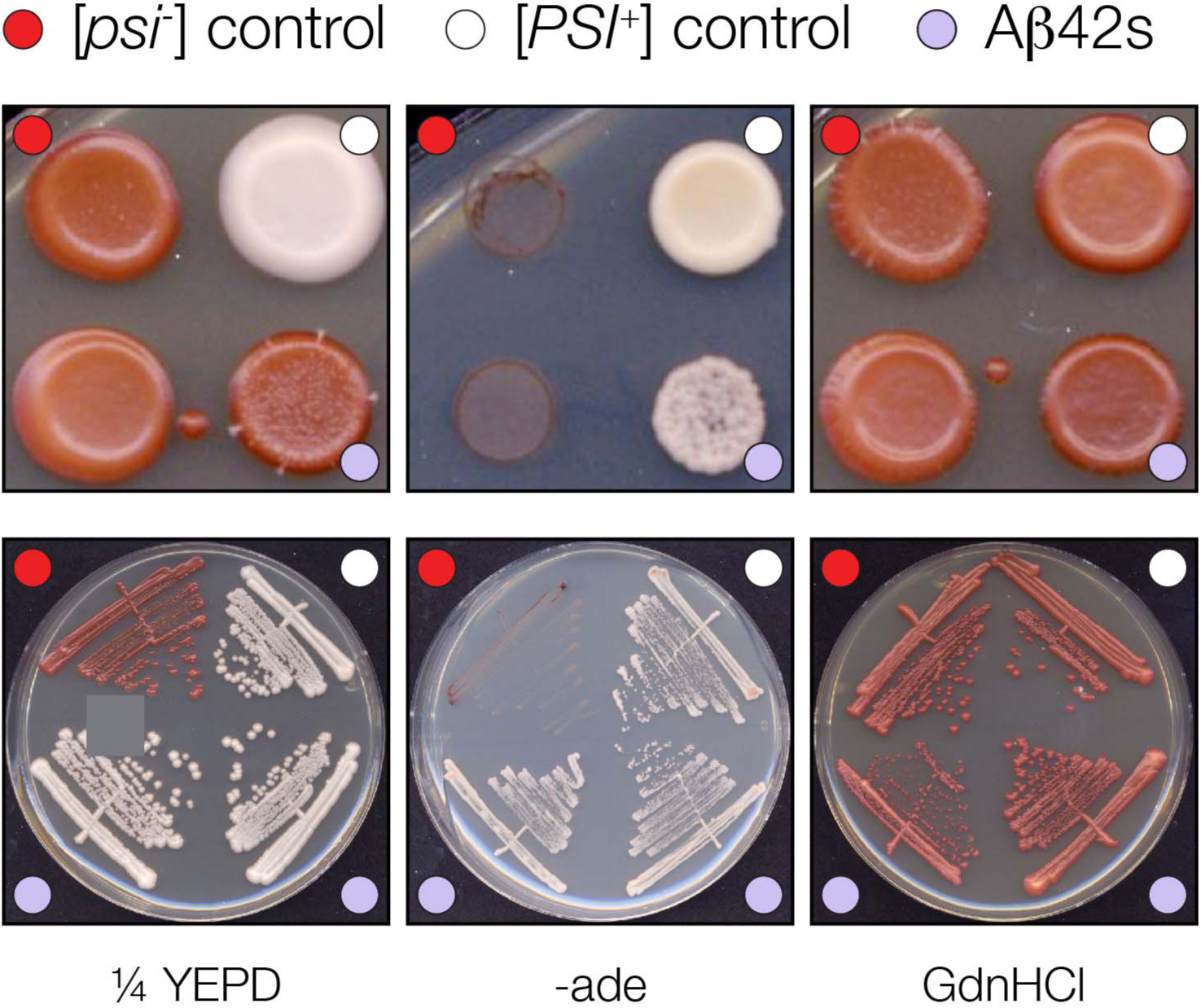
Transfection of yeast cells with Aβ42s results in enhanced [*PSI*^*+*^] conversion. A typical positively converted colony by Aβ42s is shown together with [*PSI*^*+*^] and [*psi*^*-*^] controls on replica plated as well as streaked plates. The cells were plated onto ¼ YEPD and -ade synthetic media to check for the [*PSI*^*+*^] prion phenotype and ¼ YEPD supplemented with 3mM GdnHCl to eliminate any false positives.

**Figure 8.**
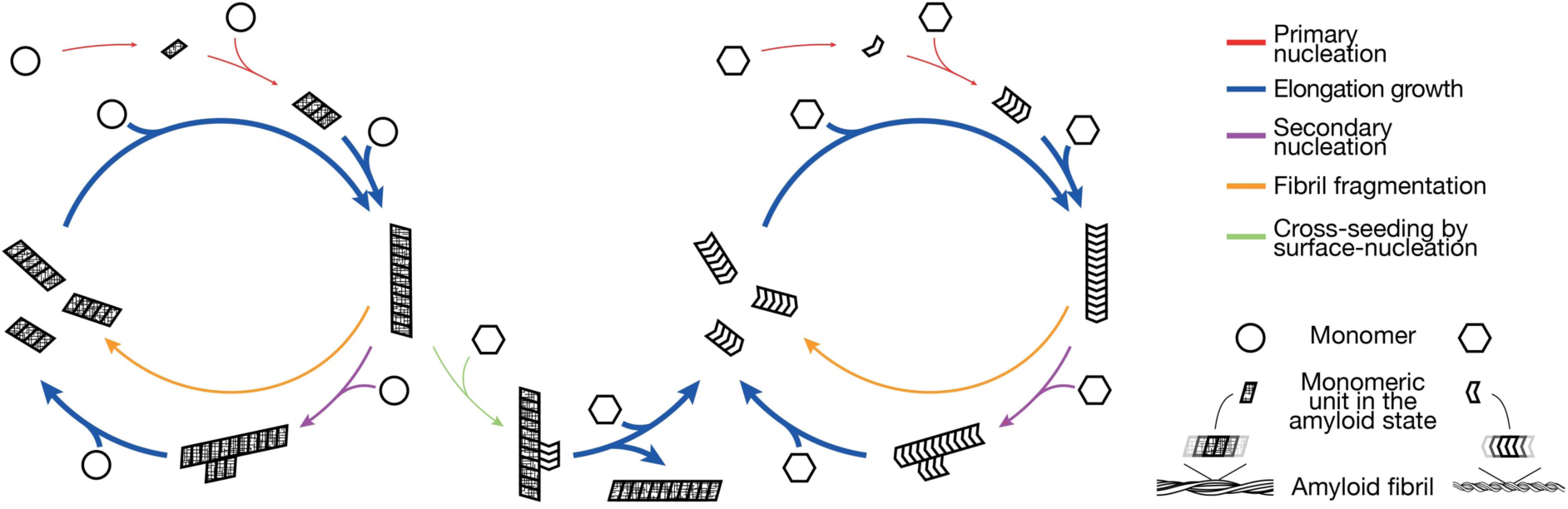
Schematic illustration of a cross-seeding mechanism involving surface-catalysed heterogeneous nucleation by amyloid particles acting as promiscuous nanoparticles with active surfaces that also promote secondary nucleation. The surface-catalyzed cross-seeding is represented by the green arrow that links the lifecycles of two otherwise unrelated amyloid systems. The arrows represent dynamic and reversible steps along the lifecycle and the thickness of the arrows illustrate typical relative magnitudes of the rates involved in each of the processes.

Similar to mammalian prion protein PrP that can exist in structurally different prion conformation and distinct “strains” (Aguzzi et al., 2007), the yeast [*PSI*^*+*^] prion linked to the amyloid form of Sup35 can also exist in different conformational variants generating phenotypically distinct “strains” (Tanaka et al., 2006). The different phenotypes reflect the strength of their biological prion activity (i.e. the characteristic stop codon read-through phenotype for the [*PSI*^*+*^] prion) and are based on the different mechanical and structural properties of the Sup35 assemblies (Frederick et al., 2014; Krishnan and Lindquist, 2005; Tanaka et al., 2006). Importantly, the prion phenotypes can be detected and visualized using a well-established colorimetric assay where *ade1-14* colonies of different [*PSI*^*+*^] strains or [*psi*^*-*^] in *Saccharomyces cerevisiae* can be identified by their colour which ranges from red to white depending on the variant (Derkatch et al., 1996; Tanaka et al., 2006; Uptain et al., 2001). Thus, the [*PSI*^*+*^] phenotype provides a sensitive *in vivo* test of prediction IV (**Figure 1 prediction IV**), i.e. whether fibrils formed from self-seeding with Sup35NMs were comparable to those formed by heterologous cross-seeded reactions through generic surface catalyzed action of Aβ42 seeds as nanoparticles. We introduced, non-seeded (i.e. *de novo* grown from Sup35NMm), self-seeded and Aβ42s cross-seeded Sup35NM particles into yeast cells by protein transfection (See Materials and Methods and (Marchante et al., 2017)). The resulting colonies were picked, stamped and grown for 7 days on three different selective plates: -ade, ¼ YEPD and ¼ YEPD+3 mM GdnHCl to score for the [*PSI*^*+*^] phenotypes induced by each of the introduced amyloid samples (**Figure 6c**). The Sup35NM particles in all three cases grown from different seeds or no seeds resulted in the same prion phenotype pattern amongst the [*PSI*^*+*^] transfectants, with the majority of [*PSI*^*+*^] colonies showing a ‘strong’ phenotype (i.e. white colonies) with a low number (around 15%) of weak [*PSI*^*+*^] colonies (i.e. pink or dark pink colonies) (**Figure 6d**). These findings further support the conclusion that the fibril particles had indistinguishable morphology independently of the seeds they were exposed to, and that they also give rise to the same biological phenotype, which in turn is sensitive to small conformational difference in the amyloid architecture (Frederick et al., 2014; Tuite et al., 2014). Therefore, these results demonstrated that seeding of Sup35NM monomers with heterologous Aβ42 amyloid seeds did not affect the conformation of Sup35 that was faithfully transmitted and propagated *in vivo*, in agreement with the structural predictions III and IV (**Figure 1 predictions III and IV**) for a cross-seeding mechanism involving generic heterogeneous nucleation catalyzed by surfaces of seeds as nanoparticles.

### Aβ42 amyloid fibril seeds are capable of inducing [*PSI*^*+*^] phenotype upon transfection into yeast cells

The results described above show that the surfaces of Aβ42s seeds can interact and cross-seed the formation of Sup35NM amyloid fibrils *in vitro* by acting as nanoparticles that promote surfaces-catalyzed interactions. To test whether heterologous Aβ42s nanoparticles introduced into a yeast cell can trigger the appearance of the Sup35-based [*PSI*^*+*^] prion as suggested by the *in vitro* experiments (**Figure 3, 4 and 5**), we next transfected yeast spheroplasts with the *in vitro* assembled Aβ42s particles as previously described (see Materials and Methods and (Marchante et al., 2017)). Strikingly, when transfected colonies were subsequently stamped onto ¼ YEPD plates, few colonies (∼2%, **Table 1**) formed with mixed white and red cells, indicative of slow conversion of the yeast cells to the [*PSI*^*+*^] phenotype compared to the cells transfected by Sup35NM fibrils. When re-streaked onto fresh selective plates, these rare colonies displayed all of the properties associated with the presence of the [*PSI*^*+*^] prion: white colonies on ¼ YEPD plates, growth on -ade plates, and red colonies on ¼ YEPD + 3 mM GdnHCl plates, the latter indicating ‘curing’ of the [*PSI*^*+*^] prion (Tuite et al., 1981). While the efficiency of transfection and subsequent conversion to [*PSI*^*+*^] was much less frequent using Aβ42s than transfection with *in vitro* formed Sup35NM particles (Marchante et al., 2017; Tanaka et al., 2006), the frequency of [*PSI*^*+*^] occurrence was around 1000-fold higher compared to spontaneous *de novo* formation of [*PSI*^*+*^] in non-transfected [*psi*^*-*^] cells (**Table 1**). Taken together, these observations show that Aβ42s particles are able to increase the appearance of the [*PSI*^*+*^] prion phenotype when introduced in yeast cells through prion transfection despite no biological or structural links existing between Sup35, its maintenance chaperone network *in vivo*, and Aβ42. In summary, these results are consistent with the promotion of non-native heterologous surface interactions by Aβ42s particles *in vivo* as we have seen *in vitro*. Taken together, the results demonstrate that that heterologous cross-seeding of amyloid may reflect the generic property of amyloid seeds and their surfaces in biological systems.

**Table 1:** Frequency of [*PSI*^*+*^] appearance for yeast cells transfected with Aβ42s compared with *de novo* formation of [*PSI*^*+*^] in wild type *Saccharomyces cerevisiae* strains.

| Strain | Frequency of $[PSI^+]$<br>appearance<sup>*</sup> / % | 95% confidence<br>interval / % |
| --- | --- | --- |
| **74D-694 transfected with Aβ42s** | $2.2 \cdot 10^{-2}$ | $1.5 \cdot 10^{-2} - 2.9 \cdot 10^{-2}$ |
| **74D-694 Haploid** | $1.1 \cdot 10^{-5}$ | $0.1 \cdot 10^{-5} - 2.1 \cdot 10^{-5}$ |
| **74D-694 Diploid** | $2.4 \cdot 10^{-5}$ | $1.5 \cdot 10^{-5} - 3.2 \cdot 10^{-5}$ |
| **243/6a<sup>†</sup>** | $10^{-7} - 10^{-5}$ | — |
| **74D-694<sup>‡</sup>** | $5.8 \cdot 10^{-7}$ | $4.6 \cdot 10^{-7} - 7.5 \cdot 10^{-7}$ |

<sup>\*</sup> Median values in $[psi^-][PIN^+]$ yeast cells; <sup>†</sup> (Lund and Cox, 1981); <sup>‡</sup> (Lancaster et al., 2010)

## DISCUSSION

Synergetic heterologous interactions of amyloid aggregates, as exemplified by amyloid cross-seeding, is a well-known phenomenon and frequently studied in relation to human diseases linked to human amyloidogenic proteins (reviewed in Sarell et al., 2013). In these cases, cross-seeding has been assumed to also contribute to why some amyloid-associated diseases coincide with the formation of other non-homologous amyloid aggregates. However, the molecular mechanism of how such cross-seeding processes proceed *in vitro* or *in vivo* remain unresolved because the widely accepted templated elongation model does not explain how completely different sequences and structures are capable of templating each other. In addition, the importance of primary sequence similarity for the observed cross-seeding between heterologous protein aggregates is not clear (e.g. Lu et al., 2020; Shida et al., 2020). Thus, the current view of cross-seeding through templated elongation alone does not readily explain the synergetic statistical links between amyloid diseases associated with non-homologous proteins (Biessels et al., 2006; Daniele et al., 2018; Sims-Robinson et al., 2010). Here, under rigorously controlled conditions, we have investigated the cross-seeding interactions between two amyloid-forming proteins, namely human Aβ42 and yeast Sup35NM. These two proteins were chosen because they are entirely unrelated in terms of sequence, structure, and biological function and consequently should not be able to template the elongation growth of each other. Yet, we observed cross-seeding effects between these two proteins, where non-homologous fibrillar seeds significantly shortened the lag phase of amyloid-forming reactions, albeit to a much lesser extent compared to homologous seeds. We also found that the effect of non-homologous cross-seeding in the fibril forming reaction for these two proteins was not symmetric, and instead depended on the aggregation properties of monomers under the reaction conditions used (Peduzzo et al., 2020), and on the specific the type of seeds used. In addition, *in vivo* studies whereby amyloid seeds can be introduced into a genetically marked strain of *S. cerevisiae* that reports the ability of those seeds to trigger the formation of Sup35 amyloid (Tanaka et al., 2006) were carried out. Whereas different *in vitro*-generated or extract-purifies variants of Sup35NM fibrils when transfected into such yeast cells are faithfully propagated, creating different ratios of weak or strong [*PSI*^+^] phenotypes (Diaz-Avalos et al., 2005; Krishnan and Lindquist, 2005; Tanaka et al., 2006), Sup35NM fibrils formed in Aβ42-seeded reaction generated a mixture of [*PSI*^+^] phenotypes that were indistinguishable to those arising when *de novo* formed or self-seeded Sup35NM fibrils generated *in vitro* were used (**Figure 6**).

The results we obtained for the heterologous seeding action between these two unrelated amyloidogenic proteins are not consistent with templated elongation, but are entirely consistent with the amyloid seeds acting as nanoparticles that affect heterologous amyloid assembly through surface-based interactions. In our experiments, both the Aβ42 and Sup35NM seeds acted to accelerate the amyloid formation of the other protein and giving rise to the observed cross-seeding effect, without acting as templates of their own conformation. Indeed, heterologous surface catalysis is often utilized in organic synthesis while amyloid seeding by surface-catalyzed heterogeneous nucleation or retardation by surface interactions are commonly observed effects of polymeric and non-polymeric nanoparticles alike (Linse et al., 2007).

Since nucleated protein assembly catalyzed by seed surfaces is not dictated by the amyloid conformation of the seeds in the same way as in templated elongation reactions, the resulting amyloid fibrils from surface-catalyzed reactions could have structures distinct from that of the seeds (**Figure 1 prediction III**). Furthermore, the resulting amyloid structures could be diverse in their morphology. This is because seeding and cross-seeding through surface-catalyzed nucleation events may be expected to introduce heterogeneity in the resulting amyloid aggregates depending on the assembly conditions, whereas seeding though templating will propagate specific amyloid conformations and thereby reducing possible heterogeneity. For amyloidogenic proteins involved in misfolded protein diseases, heterologous amyloid particles would potentially allow the formation of a pallet of conformational variants or strains, with different levels of toxicity and infectious potential (Peelaerts et al., 2015; Qiang et al., 2017). Thus, structural polymorphism (Adamcik and Mezzenga, 2018; Fändrich et al., 2018; Lutter et al., 2019) as a consequence of species heterogeneity modified by the cross-seeded reaction could lead to the generation of new toxic conformers, and their propagation could play an important role in disease.

Previous studies have demonstrated that Aβ42 aggregation is accelerated by an auto-catalysed nucleation process. In common with the cross-seeding mechanism we address here, this autocatalytic process of secondary nucleation is a surface-driven process (Cohen et al., 2013) and is one of the major process involved in homologous amyloid assembly (**Figure 1**). Thus, the surfaces of preformed amyloid seeds of Aβ42, even outside of active elongation sites at fibril ends (Milanesi et al., 2012), appear to be particularly active and capable of accelerating formation of new amyloid through promoting surface interactions. The secondary nucleation mechanism can accelerate amyloid formation as well as to generate small oligomeric species that could be biologically active in driving the toxic potential of Aβ42 amyloid (Cohen et al., 2013). This suggests that the surfaces of small Aβ42 amyloid seeds, as nanoparticle surfaces, may be able to act as general amyloid formation catalysts for both homologous and heterologous sequences, and in the process of accelerating heterologous amyloid formation generate some species that may possess cytotoxic potential. Interestingly, recent research has shown that for another yeast prion-forming protein Ure2, the surface-catalyzed secondary nucleation process does not dominate homologous assembly in presence of preformed fibrils (Yang et al., 2018). It has been suggested that the absence of secondary nucleation results in a reduced generation of toxic oligomeric species and therefore lower toxicity associated with formation and propagation of yeast prions. In contrast, a secondary nucleation mechanism has been inferred in cases involved in the amyloid formation of peptides and proteins associated with neurodegenerative diseases and type 2 diabetes (e.g. Aβ42, Aβ40, a-synuclein, IAPP, insulin; reviewed in (Linse, 2017)). Thus, the generation of such amyloidogenic species with heightened toxic potential may depend on the surface properties of the seed particles and how these surfaces interact with both homologous or heterologous monomeric protein sequences that are present in the same biological milieu.

The enhanced likelihood of generating the Sup35-based [*PSI*^*+*^] prion upon transfection of prion-free [*psi*^*-*^] cells with Aβ42 fibril particles is consistent with our hypothesis that amyloid seeds as nanoparticles can influence the formation of heterologous amyloid via aberrant surface interactions. The presence of aberrant surfaces such as those presented by Aβ42 fibril particles in yeast cells may also act through modulating cellular proteostasis. This could therefore subtly affect the balance of proteins such as the molecular chaperones critical for the prion generation and propagation pathways e.g. the AAA^+^-ATPase Hsp104 (Chernoff et al., 1995; Satpute-Krishnan et al., 2007; Shorter and Southworth, 2019). Thus, Aβ42 fibril particles *in vivo* may provide surface-mediated interactions accelerating the formation and the propagation of the amyloid state in the cells though direct as well as indirect modes of action. These insights bring to the fore surface properties and surface interactions of amyloid particles as a key mesoscopic property to target in order to understand the origin of the amyloid cytotoxic potential and the synergetic link between different amyloid diseases, as well as designing therapies to combat the disease processes associated with toxic amyloid.

## MATERIALS AND METHODS

### Protein expression and purification

Sup35NM protein samples were produced as described previously (Marchante et al., 2017) with minor changes as follows. The DNA sequence encoding the N-terminal NM region of the yeast Sup35 protein (residues 1-254) was amplified from plasmid pUKC1620 by PCR and cloned into pET15b as a *Bam*HI-*Nde*I fragment, resulting in an N-terminal His_6_-tag fusion protein. The resulting plasmid (pET15b-His_6_-NM) was then transformed into the *E. coli* strain BL21 DE3 (*F– ompT gal dcm lon hsdSB(rB-mB-) λ(DE3 [lacI lacUV5-T7 gene 1 ind1 sam7 nin5]*). For protein expression, this *E. coli* strain was grown overnight in 50 ml LB supplemented with 0.1 mg/ml ampicillin and 0.03 mg/ml chloroamphenicol, and then transferred to 1L cultures of the same medium. On reaching an OD_600_ of ∼0.5, expression was induced by the addition of IPTG (1 mM final concentration) for 4 hours. Cells were harvested at 6000 rpm and the cell pellets washed once in buffer A1 (20 mM Tris-HCl pH 8.0, 1 M NaCl, 20 mM imidazole). Cells were pelleted again, and the pellets kept at -80°C for later use. For the affinity purification step, buffer A2 (20 mM Tris-HCl pH 8.0, 1 M NaCl, 20 mM Imidazole, 6 M GdnHCl) was added to frozen cell pellets at a 5:1 (v/v) ratio, followed by sonication at an amplitude of 22 microns until the cell pellet was completely disrupted. This solution was then spun down at 13000 rpm for 30 minutes and the supernatant collected. 2 ml of Chelating Sepharose Fast Flow (GE Healthcare) was added to a small plastic column and prepared for affinity purification by sequential washing with 1 column volume (CV) of water, 0.2 M NiCl_2_, buffer A1 and buffer A2. The equilibrated resin was then resuspended in buffer A2 and added to previously collected supernatant. This mixture was then incubated for 1 hour at room temperature with agitation to improve protein binding to the affinity resin. Centrifugation at 5000 rpm was subsequently used to collect the resin, which was then washed in 5ml buffer A2, resuspended in buffer A2, and transferred back to the column. After one wash with 1 CV buffer A2, elution was achieved by addition of 3ml buffer A3 (20 mM Tris-HCl pH 8.0, 1 M NaCl, 0.25 M imidazole, 6M GdnHCl). The resulting eluate was immediately used for size-exclusion purification, which was run using a HiLoad 16/600 Superdex 200 pg (GE Healthcare) column in an ÄKTA Prime Plus chromatography system (GE Healthcare). The eluate was injected into the size-exclusion column previously equilibrated with 1 CV water followed by 1 CV buffer S1 (20 mM Tris-HCl pH 8.0, 0.5 M NaCl) and 1 CV buffer S2 (2 0mM Tris-HCl pH 8.0, 0.5 M NaCl, 6 M GdnHCl). The relevant Sup35NM protein fractions were collected according to the A_280_ displayed throughout the run, diluted to 20 µM in buffer S2 and immediately used in fibril-forming reactions.

The Amyloid Beta (1-42) peptide (Aβ42) was purchased in 5 mg batches from Bachem (Germany). This was aliquoted in 0.5 mg stock batches and frozen at -20°C. Monomers were further purified as described previously (Hellstrand et al., 2010; Walsh et al., 2009) with minor modifications. Briefly, the Aβ42 was purified using gel filtration as follows: 0.5-1 mg of Aβ42 was dissolved in 1 ml 6 M GdnHCl. This was loaded onto a Superdex 75 10/300 GL column pre-equilibrated with 2 column volumes (CV) of 20 mM sodium phosphate, pH 7.4, 0.01% NaN_3_ (buffer E). The monomer peak was eluted with buffer E and put on ice. The concentration was then determined using UV spectroscopy (280 nm) and adjusted to working concentration of 10 µM with buffer E before immediately proceeding with fibril forming reactions.

### Fibril formation and monitoring

For Sup35NM fibril formation, 2.5 ml of 20 µM purified Sup35NM were buffer exchanged into Fibril Forming Buffer (FFB - 20 mM sodium phosphate buffer pH 7.4, 50 mM NaCl) using a PD-10 column (GE Healthcare) as per the manufacturer’s instructions. Protein concentration was measured using A280 and then adjusted to 10µM using FFB. Protein samples were aliquoted into Protein LoBind tubes (Eppendorf) and polymerised at 30°C quiescently for at least 48 hours. For monitoring polymerisation, 100µl samples of protein were aliquoted into black low binding hydrograde 96-well plates (BRAND) and Thioflavin T (ThT) was added to a final concentration of 10 µM. The plate was sealed with Starseal Advanced Polyolefin Film (Starlab) and kinetics were monitored in a 96-well format (Xue et al., 2008) using a FLUOstar OMEGA plate reader (BMG Labtech) quiescently at 30°C.

For Aβ42 fibrils assembly reaction, 10 uM solution of Aβ42 monomers purified as described above, were either aliquoted either into Protein LoBind tubes or into black hydrograde 96-well plates (BRAND) with 10 µM of ThT for kinetic monitoring using identical method as described above for Sup35NM.

### Fibril fragmentation

Fibril fragmentation was achieved by sonication over different periods of time using a probe sonicator (Qsonica Q125) at 20% amplitude in consecutive 5 s on/off cycles on an ice-cooled water-bath.

### Dynamic light scattering (DLS)

All vials, tubes and cuvettes used for preparing the samples were clean dry. All solvents used were filtered to remove any particulates that may interfere with the results obtained. The Aβ42 or Sup35NM fibril seed samples obtained after controlled sonication were diluted 10 times using the same FFB as described above. The samples were subsequently characterised by DLS at 25 °C using an Anton Paar Litesizer™ 500 instrument, and the data processed using KalliopeTM Professional.

### Yeast transfection with amyloid fibrils

For yeast transfection with Sup35NM synthetic amyloid fibrils, a [*psi*^-^] derivative of the yeast strain 74D-694 (*MATα ade1-14 trp1-289 his3Δ-200 ura3-52 leu2-3,112*), and [*PIN*^*+*^] derivative of the same strain was used for transfections of Aβ42 fibrils. The transfection procedure was as previously described (Marchante et al., 2017). Briefly, cells freshly grown in YEPD to an OD_600_ of 0.5 were washed, resuspended in 12 ml ST buffer (1 M sorbitol, 10 mM Tris-HCl pH 7.5) and spheroplasts were prepared by addition of 600 U of lyticase (Sigma L4025) and 10 mM DTT during incubation at 30°C with agitation for 45 min. Spheroplasts were then harvested by centrifugation (400 *g*, 5 min), washed with 1.2M sorbitol and STC buffer (1.2 M Sorbitol, 10 mM Tris-HCl pH 7.5, 10 mM CaCl_2_) and then resuspended in 1ml STC buffer. Pre-mixture of 2 µl (approximately 1µg) of plasmid DNA (pRS416), 10µl single-stranded DNA (10 mg/ml) and 10 µl of freshly sonicated amyloid fibrils (as described above) were combined with 100 µl spheroplast suspension for each transformation reaction. This transformation mix was then incubated for 10 min at room temperature and then 0.9 ml of PEG buffer (40% PEG 4000, 10 mM TRis-HCl pH 7.5, 10mM CaCl_2_) was added to each transformation. After 30 min at room temperature, the spheroplasts were collected by centrifugation (400 *g*, 5 min), resuspended in 200 µl SOS media and added to sterile Top agar (-URA synthetic complete media with 2% agar and 1.2 M Sorbitol) being kept at 48°C, gently mixed and then poured into agar plates previously prepared using the same media. Cells were allowed to grow for 3 - 4 days at 30°C and then colonies were individually picked into 96 well plates containing YEPD. These were grown overnight at 30°C with agitation and then replica plated onto ¼ YEPD and -ade synthetic media to check for the [*PSI*^*+*^] prion phenotype and ¼ YEPD supplemented with 3mM GdnHCl to eliminate any false positives as 3 mM GdnHCl eliminates the [*PSI*^*+*^] prion (Tuite et al., 1981). Fragmented amyloid fibrils used in transfection experiments were simultaneously prepared for imaging analysis using AFM as described below. [*PSI*^*+*^] cells arising from transfections with Aβ42 fibrils typically generated colonies consisting of a mixture of white wand red cells. In such cases, the white colonies were sub-cloned on ¼ YEPD and -ade plates and re-checked for their [*PSI*^*+*^] prion phenotype on ¼ YEPD+GdnHCl plates.

### Quantification of spontaneous *de novo* appearance frequency of [*PSI*^*+*^]

The frequency of spontaneous *de novo* conversion of yeast cells from [*psi*^*-*^] to [*PSI*^*+*^] was quantified using an adaptation of a previously published protocol (Chernoff et al., 1999). Here, the [*psi*^*-*^][*PIN*^*+*^] derivative of the strain 74-D694 was used. Ten independent freshly grown colonies were randomly selected from YEPD plates and grown in YEPD media to reach an OD_600_ of 0.5. The cells were then washed, resuspended in sterile water and plated on -ade medium at 3 dilutions (around 10^5^ cells/plate, 10^4^ cells/plate and 10^3^ cells/plate) to ensure that the residual growth on -ade plates does not affect the appearance of Ade^+^ colonies. To determine the concentration of viable cells, an aliquot of each culture was used for a serial dilution plated on YEPD medium. Ade^+^ colonies were counted on -ade plates after 10 days of growth at 25°C. To confirm [*PSI*^*+*^] phenotype, the Ade^+^ colonies were replica plated on GdnHCl media and characterised further by SDD-AGE for confirming the presence of SDS-resistant Sup35 aggregates. For the Aβ42-transfected cells, colonies selected on -ura plates 3 days after transfection were grown overnight in YEPD media and stamped on –ade, ¼ YEPD and ¼ YEPD media containing 3 mM GdnHCl. [*PSI*^*+*^] formation rates were calculated according to the formula *R = f/*ln*(NR)*, where *R* is the rate of [*PSI*^*+*^] formation, *f* is the observed frequency of [*PSI*^*+*^] colonies, and *N* is the number of cells in the culture (Chernoff et al., 1999; Drake, 1991).

### Atomic Force Microscopy (AFM) analysis

The fibril samples were diluted 1:100 for Sup35NM and 20 µl droplets were deposited on freshly cleaved mica discs (Agar Scientific F7013). After 10 min incubation at room temperature, excess sample was removed by washing with 1 ml of 0.2 µm syringe-filtered mQ H_2_O and the specimens were then dried under a gentle stream of N_2_(g). For Aβ42 fibrils, samples were diluted 1:10 and 10 µl were deposited on mica disc, let dry at room temperature, washed with 500 µl of mQ H_2_O and then dried under a gentle stream of N_2_(g). Samples were imaged using a Bruker Multimode AFM with a Nanoscope V controller and a ScanAsyst probe (Silicone nitride tip with nominal tip radius = 2 nm, nominal spring constant 0.4 N/m and nominal resonant frequency 70 kHz). Images were captured at a resolution of 4.88 nm per pixel scanned. All images were processed using the Nanoscope analysis software (version 1.5, Bruker). The image baseline was flattened using 3^rd^ order baseline correction to remove tilt and bow. Processed image files were opened and analyzed using automated scripts written in Matlab (Xue, 2013).

## AUTHOR CONTRIBUTIONS

N.K. designed the research, conducted the experiments, and analyzed the data. R.M., T.J.P, and J.H. conducted the experiments, and analyzed the data. M.F.T. designed the research and analyzed the data. W.F.X. designed the research, wrote the analytical software tools, analyzed the data, and managed the research. The manuscript was written through contributions of all authors.

## ACKNOWLEDGEMENTS

We thank the members of the Xue group, and the Kent Fungal Group for helpful comments throughout the preparation of this manuscript. We thank Saskia Davis for help with AFM image analysis and Ian Brown for technical support. This work was supported by funding from the Biotechnology and Biological Sciences Research Council (BBSRC), UK grants BB/J008001/1 and BB/M02427X/1

## SUPPLEMENTARY FIGURES

**Supplementary Figure SI-1.**
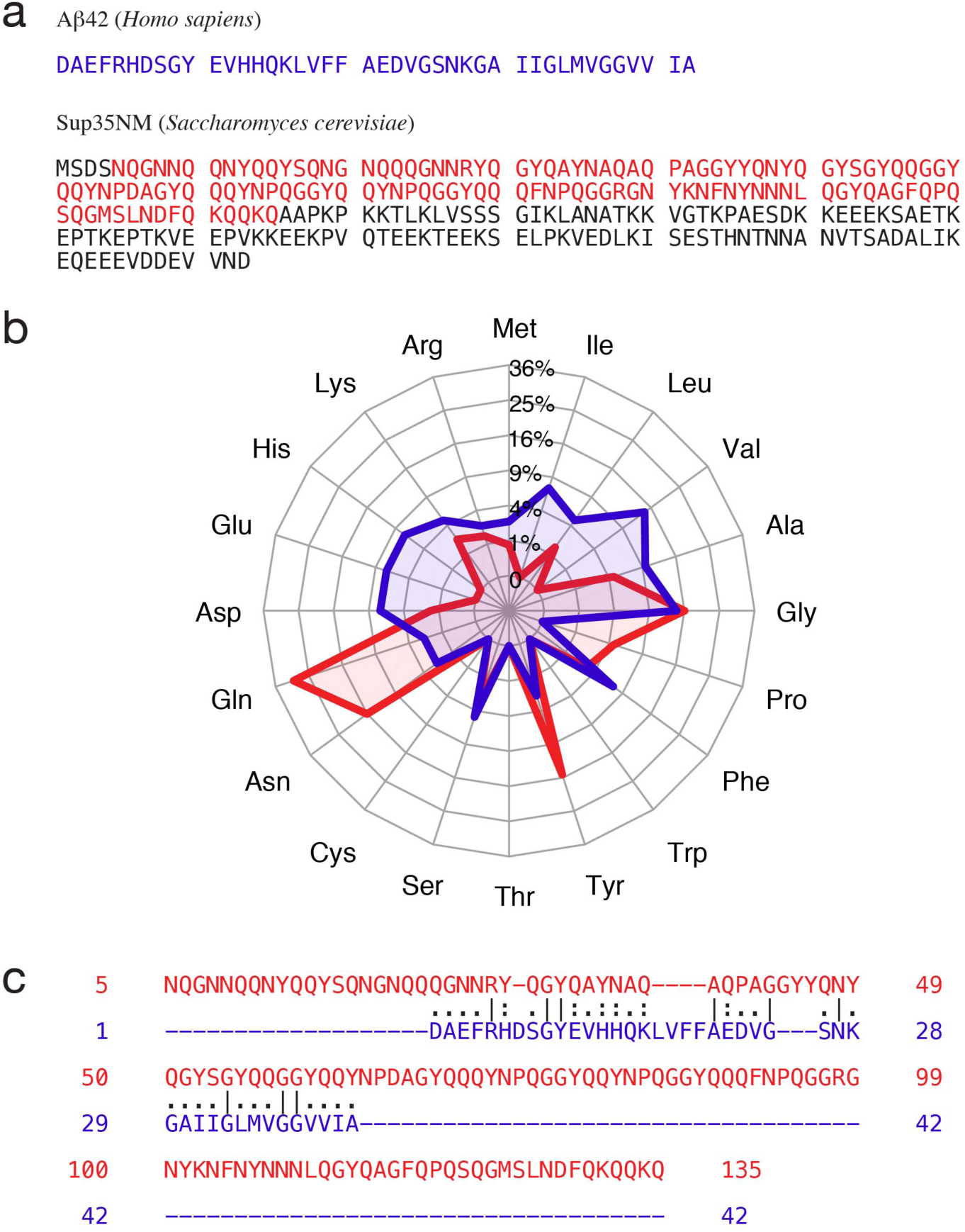
Aβ42 and Sup35NM have dissimilar amino acid compositions and low sequence similarities. (a) The primary amino acid sequences of Aβ42 and Sup35NM. For Sup35NM, the prion forming domain is highlighted in red. (b) The amino acid composition of Aβ42 and the prion domain of Sup35NM visualized in a web chart. (c) Global pairwise sequence alignment of Aβ42 and the prion domain of Sup35NM using the Needle algorithm with standard parameters. The two sequences share 6.6% identity (lines).

**Supplementary Figure SI-2.**
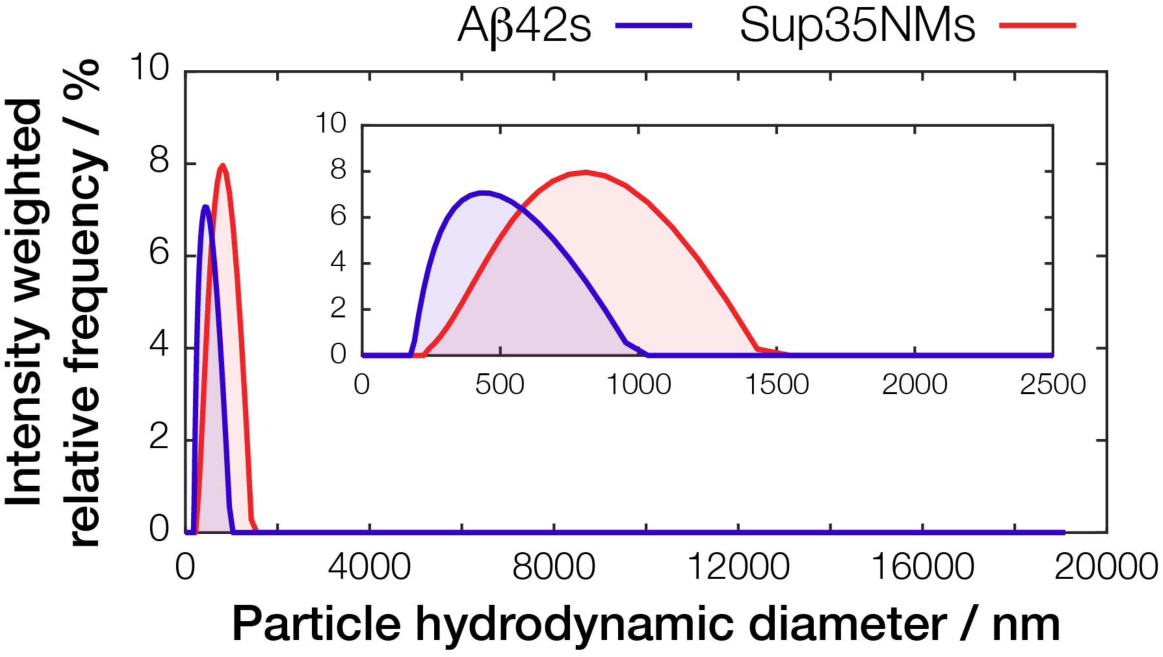
DLS characterisation of Aβ42s and Sup35NMs samples. Typical DLS traces collected at 25 °C of the same samples as those seen in **Figure 2b** at a monomer equivalent concentration of 1 µM are shown. The plot with an extended x-axis as well as an expanded view (inset) are shown for clarity.

**Supplementary Figure SI-3.**
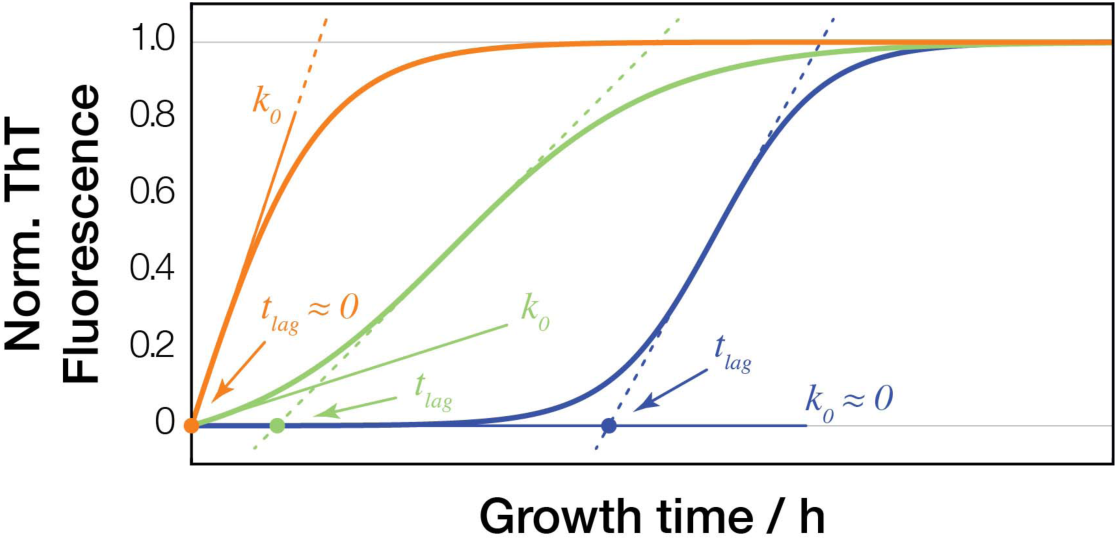
Schematic illustration of the methods used to extract *t*_*lag*_ and *k*_*0*_ values from kinetics traces of amyloid formation monitored by ThT fluorescence. Three typical example cases are shown together with indications of their *t*_*lag*_ and *k*_*0*_ values.

